# Loss of function of bHLH transcription factor Nrd1 in tomato induces an arabinogalactan protein-encoding gene and enhances resistance to *Pseudomonas syringae* pv. *tomato*

**DOI:** 10.1101/2021.11.08.467746

**Authors:** Ning Zhang, Chloe Hecht, Xuepeng Sun, Zhangjun Fei, Gregory B. Martin

## Abstract

Basic helix-loop-helix (bHLH) transcription factors constitute a superfamily in eukaryotes but their roles in plant immunity remain largely uncharacterized. We found that the transcript abundance in tomato leaves of one bHLH transcription factor-encoding gene, *Nrd1* (<u>n</u>egative <u>r</u>egulator of resistance to <u>D</u>C3000 <u>1</u>), was significantly increased after treatment with the immunity-inducing flgII-28 peptide. Plants carrying a loss-of-function mutation in *Nrd1* (Δnrd1) showed enhanced resistance to *Pseudomonas syringae* pv. *tomato* (*Pst*) DC3000 although early pattern-triggered immunity responses such as generation of reactive oxygen species and activation of mitogen-activated protein kinases after treatment with flagellin-derived flg22 and flgII-28 peptides were unaltered compared to wild-type plants. An RNA-Seq analysis identified a gene, *Agp1*, whose expression is strongly suppressed in an *Nrd1*-dependent manner. *Agp1* encodes an arabinogalactan protein and overexpression of the *Agp1* gene in *Nicotiana benthamiana* led to ∼10-fold less *Pst* growth compared to the control. These results suggest that the Nrd1 protein promotes tomato susceptibility to *Pst* by suppressing the defense gene *Agp1*. RNA-Seq also revealed that loss of Nrd1 function has no effect on the transcript abundance of immunity-associated genes including *Bti9, Core, Fls2, Fls3* and *Wak1* upon *Pst* inoculation, suggesting that the enhanced immunity observed in the Δnrd1 mutants is due to the activation of key PRR signaling components as well as loss of Nrd1-regulated suppression of *Agp1*.

## Introduction

Plants have evolved sophisticated surveillance mechanisms to rapidly recognize and respond to pathogen attacks (Lolle et al., 2020; Zhou and Zhang, 2020). The first layer of plant immunity, referred as pattern-triggered immunity (PTI), is triggered when plant cells detect microbe-associated molecular patterns (MAMPs) through transmembrane pattern recognition receptors (PRRs) (DeFalco and Zipfel, 2021). Successful pathogens deploy effectors into plant cells that interfere with PTI, leading to effector-triggered susceptibility (ETS) (Abramovitch et al., 2006). To defeat ETS, plants activate a more robust immune response, the effector-triggered immunity (ETI), where the nucleotide-binding leucine-rich repeat (NB-LRR or NLR) proteins directly or indirectly recognize a given effector, resulting in a hypersensitive cell death response (HR) and disease resistance (Jones and Dangl, 2006; Lolle et al., 2020). Although PRR-mediated PTI and NLR-mediated ETI involve different activation mechanisms and different early signaling components, recent evidence suggests that the two layers share some downstream components and both are needed to ensure robust immunity (Ngou et al., 2021; Yuan et al., 2021a; Yuan et al., 2021b)

The interaction of tomato (*Solanum lycopersicum*) with the bacterial pathogen *Pseudomonas syringae* pv. *tomato* (*Pst*) is a well-developed model system for understanding the molecular basis of plant immunity and bacterial pathogenesis (Martin, 2012; Roberts et al., 2019; Wu and Kamoun, 2019; Xin et al., 2018). When *Pst* enters the apoplastic space of the tomato leaves, two flagellin-derived MAMPs, flg22 and flgII-28, are recognized by the tomato PRRs Fls2 and Fls3, respectively (Hind et al., 2016; Roberts et al., 2020; Zhang et al., 2020). MAMP detection activates early PTI responses such as production of reactive oxygen species (ROS), activation of the mitogen-activated protein kinase (MAPK) cascades, and transcriptional reprogramming of a subset of defense genes (Jia and Martin, 1999; Li et al., 2016; Nguyen et al., 2010; Zipfel, 2014). The two *Pst* effector proteins, AvrPto and AvrPtoB, bind and interfere with the protein kinase domain of Fls2, Fls3 and the co-receptor Bak1 thus disrupting the host response to these MAMPs (Cheng et al., 2011; Hind et al., 2016; Xiang et al., 2008). The two effectors are also recognized by the host kinases Pto and Fen and trigger the hypersensitive response through the NLR protein Prf (Kim et al., 2002; Oh and Martin, 2011; Pedley and Martin, 2003).

RNA-Seq analyses have been used to identify PTI- and ETI-specific genes in the tomato-*Pst* system by inoculating plants with *Pst* strains eliciting only the PTI or ETI response (Pombo et al., 2014; Rosli et al., 2013). A subset of FIRE (flagellin-induced, repressed by effectors) genes were identified and the cell wall-associated kinase, SlWak1, was demonstrated to play a critical role in the PTI signaling pathway (Rosli et al., 2013; Zhang et al., 2020). Similarly, a subset of ETI-specific genes whose expression was induced specifically during ETI were identified and one kinase, Epk1, was shown to play a role in the host response to three effector proteins (Pombo et al., 2014). These RNA-Seq data provide a powerful resource for identifying novel immunity-associated genes involved in the tomato-*Pst* interaction.

We have recently reported the generation of hundreds of CRISPR/Cas-mediated tomato lines carrying mutations in putative immunity-associated genes (Jacobs et al., 2017; Zhang et al., 2020; Zheng et al., 2019). The availability of these tomato mutant lines provides a robust resource for the research community to test the function of specific genes in plant immunity and other biological processes (Roberts et al., 2020; Zhang et al., 2020; Zheng et al., 2019). We initially screened homozygous mutant plants by inoculating them with various *Pst* strains, including DC3000, to determine if they play a demonstrable role in PTI or ETI. Additional experimental methods including a ROS assay, MAPK activation assay, reporter gene assay, and HR assay were also applied to the mutant collection to identify new components of response pathways during the tomato-*Pst* interaction.

The basic helix-loop-helix (bHLH) proteins are a superfamily of transcription factors (TFs) that play an essential role in diverse biological processes in animals and plants (Heim et al., 2003; Li et al., 2006; Sun et al., 2015; Toledo-Ortiz et al., 2003; Wang et al., 2015a; Wang et al., 2015b). The bHLH family is defined by the bHLH signature domain, which consists of a N-terminal basic region functioning as a DNA-binding motif recognizing the E-box element (CANNTG), and a C-terminal HLH region acting as a dimerization domain to form homodimer or heterodimer required for transcription factor functions (Toledo-Ortiz et al., 2003). The bHLH TFs can transcriptionally activate or suppress target genes by specifically binding to their promoters (Hu et al., 2020; Hussain et al., 2021; Xu et al., 2014). In tomato, ∼160 bHLH protein-encoding genes were identified (Sun et al., 2015; Wang et al., 2015b), but only a few have been functionally characterized (Du et al., 2015; Kim and Mudgett, 2019; Ling et al., 2002; Schwartz et al., 2017) and even fewer have been reported to play a critical role in plant immunity (Kim and Mudgett, 2019; Schwartz et al., 2017).

The transcript abundance of one gene, encoding a bHLH transcription factor, referred to now as *SlNrd1* (*<u>S</u>. lycoperscicum* <u>n</u>egative <u>r</u>egulator of resistance to <u>D</u>C3000 <u>1</u>, hereafter *Nrd1*), was previously found to be increased in tomato leaves specifically upon treatment with flgII-28. Here, through loss-of-function analyses we found that, unexpectedly, Nrd1 appears to act as a negative regulator in tomato immunity to *Pseudomonas syringae* pv. *tomato* DC3000. Using the CRISPR-generated Δnrd1 mutant plants and RNA-Seq we identified a gene encoding an arabinogalactan protein (*Agp1*), whose expression was strongly suppressed by Nrd1. Overexpression of *Agp1* in *Nicotiana benthamiana* led to significantly less *Pst* growth, indicating it is a Nrd1-regulated defense gene against *Pseudomonas syringae*.

## Results

### Identification of Nrd1 and generation of stable loss-of-function mutants in tomato

Previous RNA-Seq analyses revealed that the transcript abundance of tomato *Nrd1* gene (*Solyc03g114230*) was significantly increased in leaves after treatment with 1 µM flgII-28 (Rosli et al., 2013), suggesting it might play an important role in the tomato-*Pst* PTI response. To study the possible role of *Nrd1* in tomato immunity, we generated three T0 knockout mutant lines in tomato cultivar RG-PtoR using CRISPR/Cas9 with a guide RNA (5’-GTAGTCCAGAAAAGCTAGAC-3’; **Fig. 1A**), which targets the first exon of the *Nrd1* gene. Two *Nrd1* independent homozygous mutants (Δnrd1-1 and Δnrd1-2) were derived and used in this study. The Δnrd1-1 mutant has a 2-bp deletion, resulting in a premature stop codon at the 27^th^ amino acid (aa) of the Nrd1 protein, whereas Δnrd1-2 contains a 13-bp deletion, causing a premature stop codon at the 18^th^ aa (**Fig. 1A**). No morphological defects were observed in either of the two *Nrd1* mutant plants when grown under greenhouse conditions (**Fig. 1B**).

**Figure 1.**
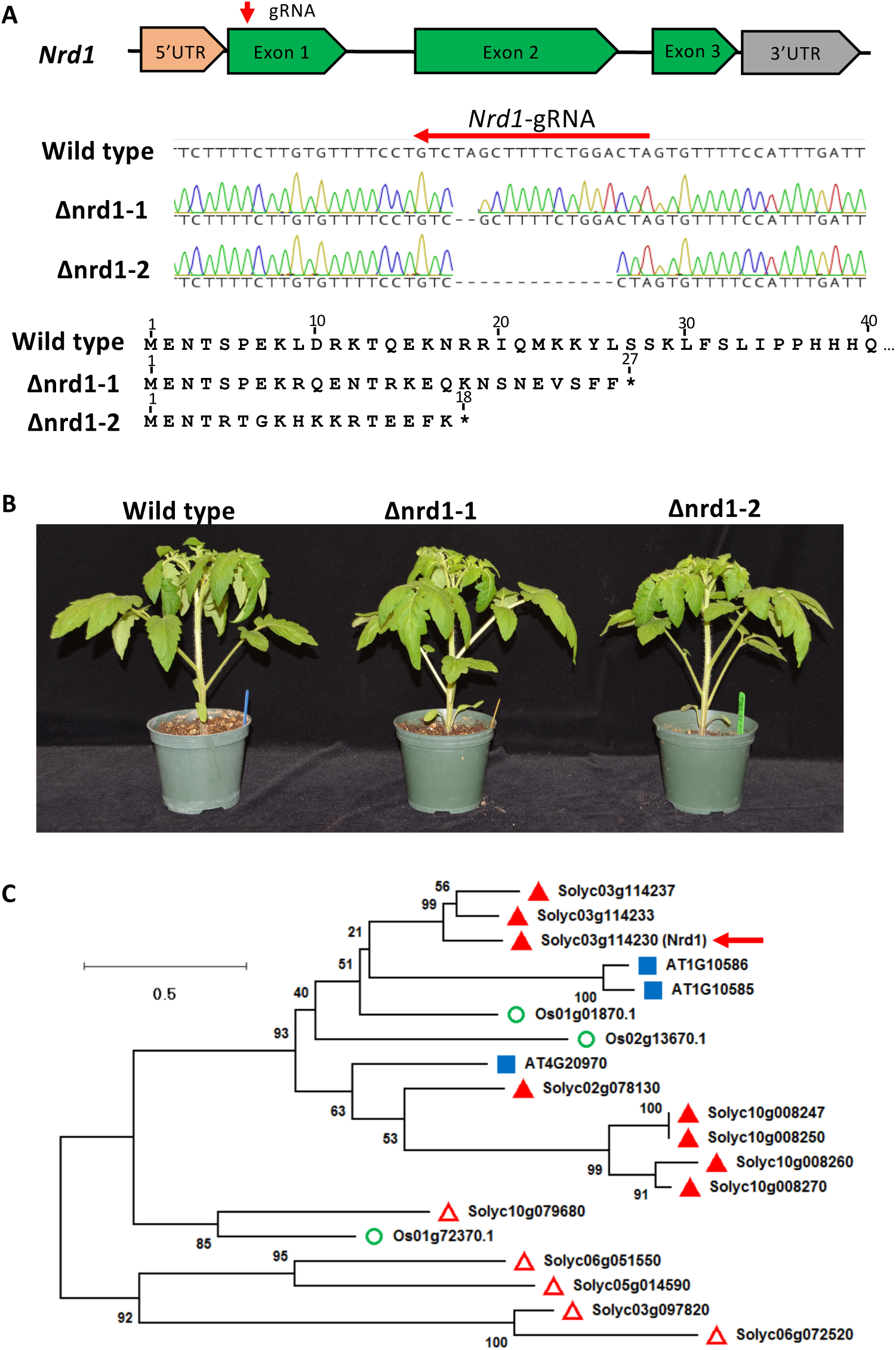
Generation of tomato Δnrd1 mutants by CRISPR/Cas9. **A**, Schematics showing the guide-RNA (gRNA) target site and the missense mutations present in two independent Δnrd1 lines. The Δnrd1-1 line has a 2-bp deletion and the Δnrd1-2 line has a 13-bp deletion. Wild type is RG-PtoR. Mutations in the lines introduce a premature stop codon at the 27^th^ or 18^th^ amino acid of the Nrd1 protein, respectively. **B**, Photographs of four-week-old wild-type RG-PtoR and the two Δnrd1 mutant lines grown in the greenhouse. **C**, Phylogenetic tree of Nrd1 and related proteins. Amino acid sequences of Nrd1 and related proteins in *Arabidopsis*, rice and tomato were used to generate a maximum likelihood tree. The tree is drawn to scale, with branch lengths measured in the number of substitutions per site. Numbers on branches indicate bootstrap support of the nodes (%). The red arrow indicates the Nrd1 protein. Proteins identified by BLAST: closed red triangle (from tomato); blue box (from *Arabidopsis*); open circle (from rice). Open red triangles indicate tomato bHLH proteins that have been characterized previously (Du et al., 2015; Kim and Mudgett, 2019; Ling et al., 2002; Schwartz et al., 2017).

*Nrd1* encodes a bHLH transcription factor containing a domain that binds the E-box motif (CANNTG) in the promoter sequence of target genes (Sun et al., 2015). To determine if *Nrd1* has close homologs in tomato, *Arabidopsis*, or rice, we performed multiple BLAST (Basic Local Alignment Search Tool) searches of the NCBI databases using the Nrd1 protein sequence as the query sequence and obtained a limited number of protein hits. Phylogenetic analysis revealed that the Nrd1 protein has two relatively close paralogs in tomato, Solyc03g114233 and Solyc03g114237 (**Fig. 1C** and **Supplemental Fig. 1A**), with 60.3% and 65.0% similarity to the Nrd1 protein sequence, respectively. Nothing appears to be known about the biological functions of the two Nrd1 paralogs, and they are newly annotated genes in the latest version of tomato reference genome (SL4.0; https://solgenomics.net). However, our RNA-Seq data revealed very low transcript levels of *Solyc03g114233* and *Solyc03g114237* in leaves of both wild-type RG-PtoR plants and Δnrd1 mutants, whereas *Nrd1* showed a much higher transcript abundance after *Pst* inoculation (**Supplemental Fig. 1B**). These results suggested that Nrd1, but not the two close paralogs, might play a role in the plant response to *Pst*. No clear orthologs of Nrd1 occur in *Arabidopsis* or rice, with the most closely related proteins (AT1G10585, AT1G10586, and Os01g01870) having a very low sequence similarity (28.3%, 29.3%, and 38.3%, respectively) to Nrd1.

### Mutations in *Nrd1* cause enhanced resistance to *Pst* in tomato

To test whether loss-of-function mutations in *Nrd1* affect the ETI response to *Pst*, we vacuum-infiltrated *Pst* DC3000 into the two Δnrd1 mutants, wild-type RG-PtoR (which expresses the *Pto* and *Prf* genes allowing recognition of effectors AvrPto/AvrPtoB; (Martin, 2012)) and RG-prf3 (which has a mutation in *Prf* that makes the Pto pathway nonfunctional) plants (**Fig. 2A**). We observed no significant difference in bacterial populations between the Δnrd1 mutants and wild-type RG-PtoR two days after inoculation, whereas bacterial populations were 10-fold more in RG-prf3 compared to Δnrd1 and RG-PtoR plants. Similarly, the Δnrd1 mutants and RG-PtoR plants had no disease symptoms whereas RG-prf3 showed severe disease symptoms six days after inoculation. These data indicate that Nrd1 does not have a major role in the ETI pathway acting against *Pst* DC3000.

**Figure 2.**
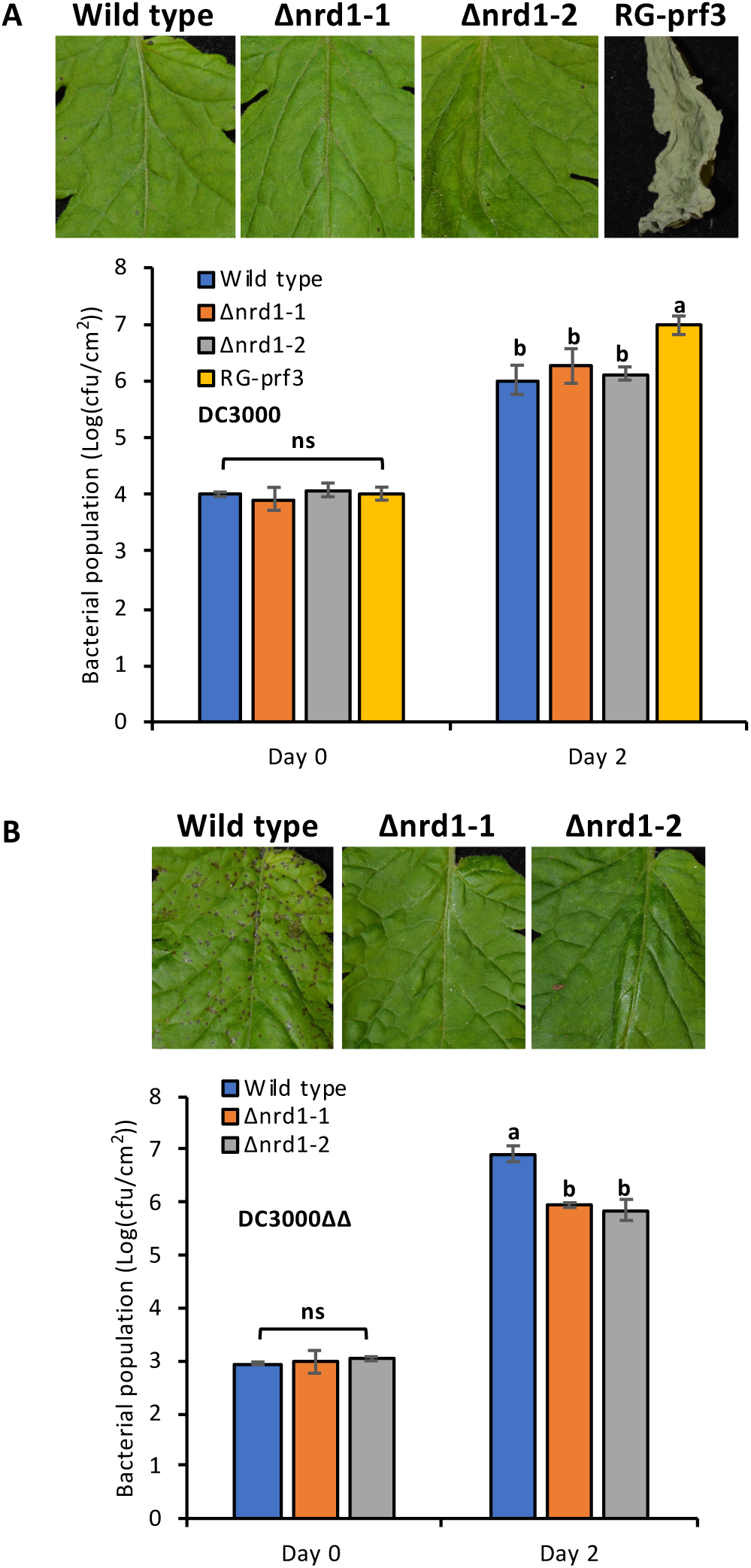
Investigation of ETI- and PTI-mediated immunity in the Δnrd1 mutants. Four-week-old Δnrd1 plants, RG-PtoR (wild type), and RG-prf3 (a *Prf* mutant) plants were vacuum-infiltrated with: **A**, 1 × 10^6^ cfu/mL DC3000 or **B**, 5 × 10^4^ cfu/mL DC3000Δ*avrPto*Δ*avrPtoB* (DC3000ΔΔ). Photographs of disease symptoms were taken at 6 days (**A**) or 5 days (**B**) after inoculation. Bacterial populations were measured at 3 hours (Day 0) and two days (Day 2) after infiltration. Bars show means ± standard deviation (SD). Different letters indicate significant differences based on a one-way ANOVA followed by Student’s *t* test (p < 0.05). ns, no significant difference. Three or four plants for each genotype were tested per experiment. The experiment was performed three times with similar results.

To test whether Nrd1 contributes to PTI acting against *Pst*, we vacuum-infiltrated the two Δnrd1 mutants and RG-PtoR with DC3000Δ*avrPto*Δ*avrPtoB* (DC3000ΔΔ) (**Fig. 2B**), which lacks the AvrPto and AvrPtoB effectors and therefore cannot activate ETI. Both mutant lines, Δnrd1-1 and Δnrd1-2, showed ∼10-fold smaller populations of *Pst* compared to wild-type RG-PtoR two days after bacterial inoculation. In addition, the Δnrd1 mutants developed much less symptoms of bacterial speck disease on leaves compared to RG-PtoR five days after inoculation. Thus, Nrd1 appears to act as a negative regulator of PTI against *Pst* DC3000, which was unexpected given that *Nrd1* transcripts increase in abundance upon treatment with flgII-28, a MAMP, and we suspected it might make a positive contribution to PTI. The enhanced resistance in the Δnrd1 mutants to DC3000Δ*avrPto*Δ*avrPtoB* was not observed in experiments with four other *Pst* strains or with *Xanthomonas campestris* pv. *vesicatoria* (also known as *X. euvesicatoria*; **Supplemental Table 1**).

### Mutations of *Nrd1* do not affect MAMP-induced ROS production or MAPK activation

ROS production and MAPK activation are two early PTI-associated responses in bacterial-inoculated plants. To investigate whether *Nrd1* contributes to these PTI responses, we performed ROS and MAPK activation assays using the two flagellin-derived peptides, flg22 and flgII-28 (**Fig. 3**). We observed no difference in either ROS production or MAPK activation in the Δnrd1-1 and Δnrd1-2 mutant lines compared to wild-type plants when treated with these peptides, indicating that role of *Nrd1* in PTI is downstream or independent of ROS and MAPK signaling pathways.

**Figure 3.**
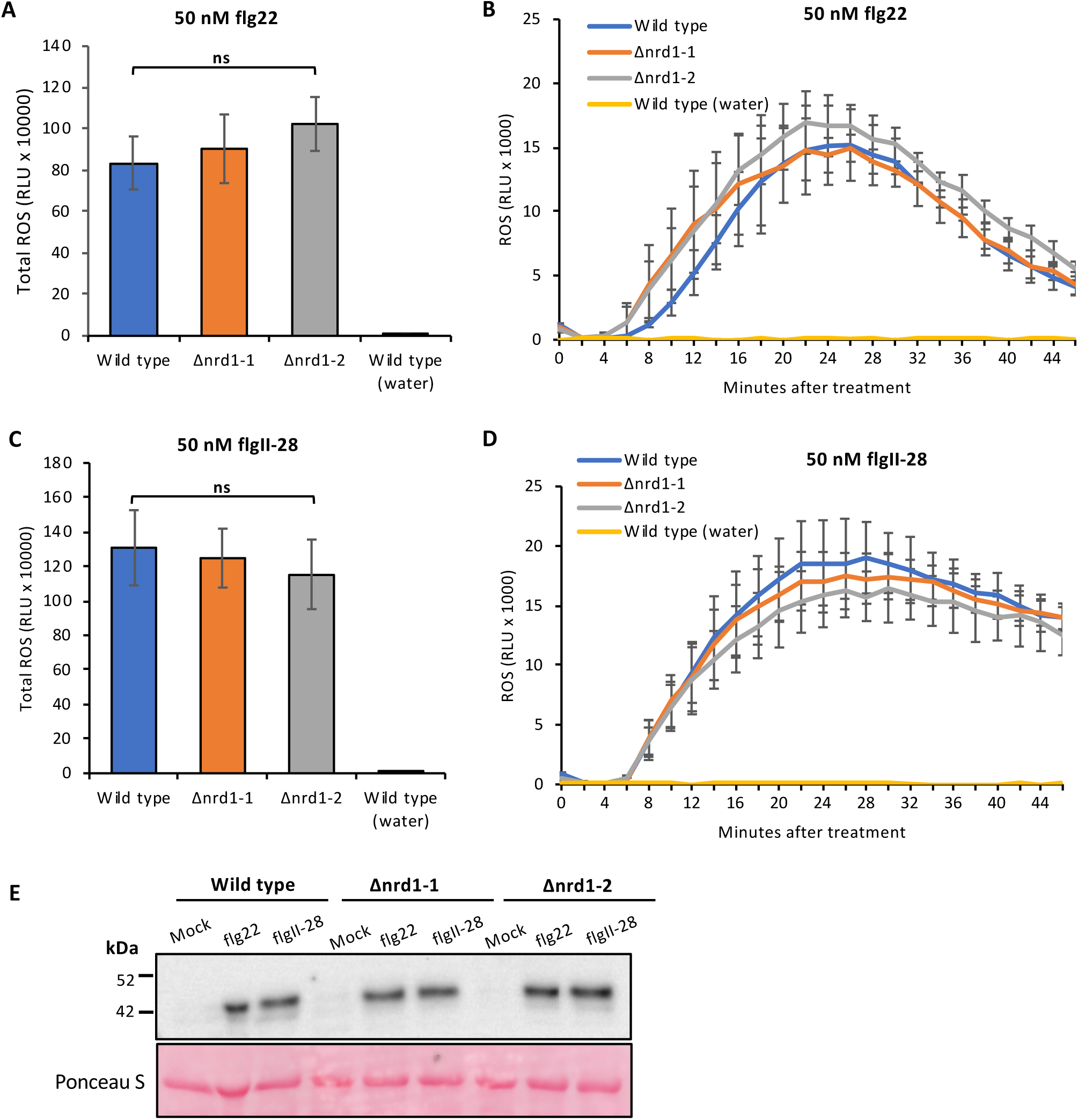
Investigation of MAMP-induced ROS production and MAPK activation in the Δnrd1 mutants. **A**-**D**, Leaf discs from Δnrd1 or RG-PtoR wild-type plants were treated with 50 nM flg22 (**A-B**), 50 nM flgII-28 (**C-D**), or water only. Relative light units (RLU) were measured over 45 minutes. One-way ANOVA followed by Student’s *t* test (p < 0.05) was performed for total ROS (**A, C**) or at 24 min (peak readout) and 45 min after treatment with flg22 or flgII-28 (**B, D**). Bars represent means ± SD. No significant difference was observed between Δnrd1 and wild-type plants in either treatment. **E**, Leaf discs from wild-type RG-PtoR plants and Δnrd1 mutants were treated with 10 nM flg22, 25 nM flgII-28, or water (mock) for 10 min. Proteins were extracted from a pool of discs from plants of the three genotypes and subjected to immunoblotting using an *anti*-pMAPK antibody that detects phosphorylated MAPKs. Ponceau staining shows equal loading of proteins.

### RNA sequencing identifies putative *Nrd1*-regulated defense and susceptibility genes

Based on the enhanced resistance to *Pst* in the Δnrd1 mutants, we hypothesize that the increased abundance of the *Nrd1* transcripts after flgII-28 treatment leads to increased Nrd1 protein that acts to suppress a subset of defense-related (D) genes and/or induces a subset of susceptibility (S) genes, thus promoting the growth of *Pst*. If this were the case, then in the Δnrd1 mutants, the Nrd1-regulated defense genes would be induced or no longer suppressed while the S genes would be suppressed, resulting in enhanced resistance to *Pst* infection. To identify possible Nrd1-regulated genes, we performed an RNA-Seq analysis using the two Δnrd1 mutants and wild-type RG-PtoR plants inoculated with DC3000Δ*avrPto*Δ*avrPtoB* (**Table 1**). Transcript levels were quantified as fragments per kilobase of transcript per million mapped fragments (FPKM), and ranged from 0 to approximately 10,000 for the genes predicted in the tomato genome. A total of 51 genes were differentially expressed in both Δnrd1-1 and Δnrd1-2 mutants compared to wild-type plants (**Supplemental Table 2**). From these, we selected six putative defense-related genes (fold-change ≥ 2 and adjusted *p* <0.05) and three putative susceptibility genes (fold-change < 0.5 and adjusted *p* <0.05), based on two criteria: 1) the transcript abundance was Δ 2 FPKM in either Δnrd1 mutants or wild-type plants; and 2) the expression of putative *Nrd1*-regulated defense genes (up-regulated in Δnrd1 mutants) was suppressed after flgII-28 treatment in wild-type plants, while the putative susceptibility (S) genes (down-regulated in Δnrd1 mutants) were induced by flgII-28 in wild-type plants, based on previous RNA-Seq data (Rosli et al., 2013) (**Supplemental Table 2**). Using the motif-searching database PlantPan2.0 (Chow et al., 2016), we found 1 to 5 copies of the E-box element (CANNTG) in the promoters of these nine candidate genes (**Supplemental Fig. 2**), indicating Nrd1 potentially binds to their promoters to either induce (S genes) or suppress (D genes) their expression.

**Table 1.** Nrd1-regulated putative defense-related genes and susceptibility genes identified by RNA-Seq. **A**, Summary of genes with increased or decreased transcript abundance in the Δnrd1 lines compared to wild type (RG-PtoR) 6 hours after inoculation with 5 × 10^6^ cfu/mL DC3000Δ*avrPto*Δ*avrPtoB* (DC3000ΔΔ). A ≥2-fold difference and adjusted p < 0.05 were used as cut-offs. **B**, Selected putative defense-related genes and susceptibility genes. *Fold change of gene expression in Δnrd1 lines compared to that in wild-type plants.

| **A** |  | **Comparison** | **Total # of differentially expressed genes** | **# of up-regulated genes** | **# of down-regulated genes** |
| --- | --- | --- | --- | --- | --- |
| | | Δnrd1-1/wild type | 463 | 211 | 252 |
| | | Δnrd1-2/wild type | 144 | 93 | 51 |
|  |  | Common | 51 | 43 | 8 |

| **B** |  | **Gene ID** | **Description** | **Δnrd1-1/wild type\*** | **Adjusted P** | **Δnrd1-2/wild type\*** | **Adjusted P** |
| --- | --- | --- | --- | --- | --- | --- | --- |
| **Putative defense-related gene (up-regulated in nrd1 mutants)** | D1 | Solyc05g024190 | Chlorophyll synthase, chloroplastic | 2.857 | 0.001313 | 3.846 | 0.00792 |
|  | D2 | Solyc07g061790 | Heme-binding protein 2-like | 4.348 | 2.45E-05 | 6.667 | 2.66E-08 |
|  | D3 | Solyc02g077330 | GDSL esterase/lipase | 2.778 | 0.014041 | 3.571 | 0.01104 |
|  | D4 | Solyc12g009650 | SI proline-rich protein | 2.222 | 6.79E-13 | 2.326 | 2.97E-05 |
|  | D5 | Solyc11g019910 | Plant invertase/pectin methylesterase inhibitor superfamily protein | 2.128 | 0.00512 | 2.632 | 5.81E-05 |
|  | D6 | Solyc08g078020 | Arabinogalactan | 3.125 | 3.81E-11 | 3.704 | 3.40E-09 |
| **Putative susceptibility genes (down-regulated in nrd1 mutants)** | S1 | Solyc03g112030 | Cytochrome P450 | 0.312 | 0.023826 | 0.415 | 0.00779 |
|  | S2 | Solyc02g088210 | SPX domain-containing protein 4 | 0.457 | 0.000645 | 0.355 | 5.14E-09 |
|  | S3 | Solyc05g007440 | ARM repeat superfamily protein | 0.289 | 3.05E-30 | 0.348 | 4.42E-19 |

### Overexpression of *Agp1* in *Nicotiana benthamiana* significantly inhibits bacterial growth

To test the possible functions of the Nrd1-regulated genes in defense or susceptibility, we performed an agromonas assay (Buscaill et al., 2021). In this assay, agroinfiltration is used first to overexpress the gene of interest in *N. benthamiana* leaves followed 2 days later by syringe-inoculation of the *Pst* strain DC3000Δ*avrPto*Δ*avrPtoB*Δ*hopQ1-1* or DC3000Δ*hopQ1-1* at the same agroinfiltrated spots (**Fig. 4** and **Supplemental Fig. 3**). HopQ is recognized by NLR Roq in *N. benthamiana* and its deletion makes DC3000 virulent on this species (Schultink et al., 2017; Wei et al., 2007)). We hypothesized that overexpression of an important defense gene would inhibit *Pst* growth, while overexpression of an essential S gene would promote *Pst* growth. Among the nine candidate genes tested, overexpression in *N. benthamiana* leaves of the putative defense-related gene D6, *Agp1* (Solyc08g078020), encoding an arabinogalactan protein, led to 8-to 10-fold less bacterial growth when inoculated with DC3000Δ*avrPto*Δ*avrPtoB*Δ*hopQ1-1* or DC3000Δ*hopQ1-1*, indicating Agp1 plays a critical role in tomato resistance to *Pst*. Expression of all proteins was confirmed by Western blot (**Supplemental Fig. 3**)

**Figure 4.**
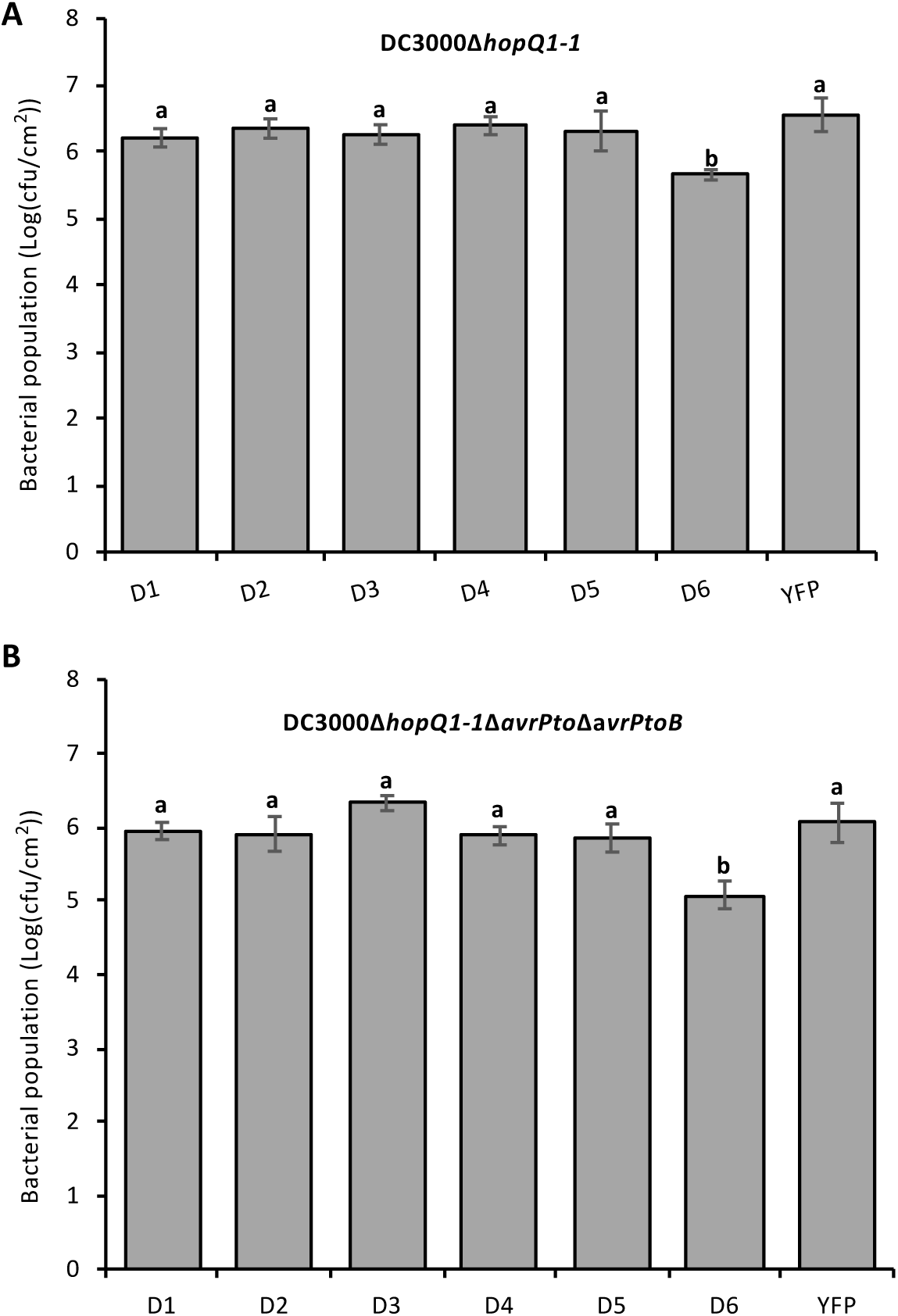
Testing putative defense-related genes in an agromonas assay. **A**-**B**, Leaves of five-week-old *Nicotiana benthamiana* plants were syringe-infiltrated with *Agrobacterium* (1D1249) strains (OD = 0.5) carrying a binary expression vector expressing each gene. Two days later, the same agroinfiltrated spots were syringe-infiltrated with 5 × 10^4^ cfu/mL DC3000*ΔhopQ1-1* or 5 × 10^4^ cfu/mL DC3000*ΔhopQ1-1*Δ*avrPto*Δ*avrPtoB* (**B**). *Pst* DC3000 populations were measured two days after the second infiltration. Bars show means ± SD. Different letters indicate significant differences based on a one-way ANOVA followed by Student’s *t* test (p < 0.05). ns, no significant difference. Three or four plants were tested with each gene in each experiment. Each experiment was performed at least two times with similar results.

Agp1 has a predicted signal peptide (SP) and a glycosylphosphatidylinositol (GPI) lipid anchor and, like other arabinogalactan proteins, it is likely associated with the outer leaflet of the plasma membrane (Silva et al., 2020). To investigate the potential function of the Agp1 SP and GPI anchor in immunity, we introduced amino acid substitutions into the SP sequence (SP-L12H and SP-T20K/A22H) or GPI-anchor sequence (GPI-S128K/S129K and GPI-F151K/F152K), or deleted the entire SP (ΔSP) or GPI-anchor sequence (ΔGPI) (**Fig. 5**). We then performed the agromonas assay to test whether the effect of these substitutions on Agp1-mediated immunity to *Pst*. All of the substitutions, except SP-L12H and GPI-S128K/S129K, impacted the ability of Agp1 to suppress *Pst* DC3000 growth compared to the wild-type Agp1 which, as expected, significantly inhibited bacterial growth in this assay (**Fig. 5**). Each of the variant proteins was expressed similar to wild-type Agp1, except for the one lacking the entire SP protein, probably due to protein degradation (**Fig. 5**). The mass of the Agp1 protein and its variants was more than twice than expected based solely on their amino acid sequences, likely due to glycosylation, as Agp1 contains 28 predicted glycosylated sites (Steentoft et al., 2013) (**Supplemental Fig. 4**). Overall, these results showed signal peptide sequence and GPI-anchor sequence are essential for Agp1-mediated resistance to *Pst*.

**Figure 5.**
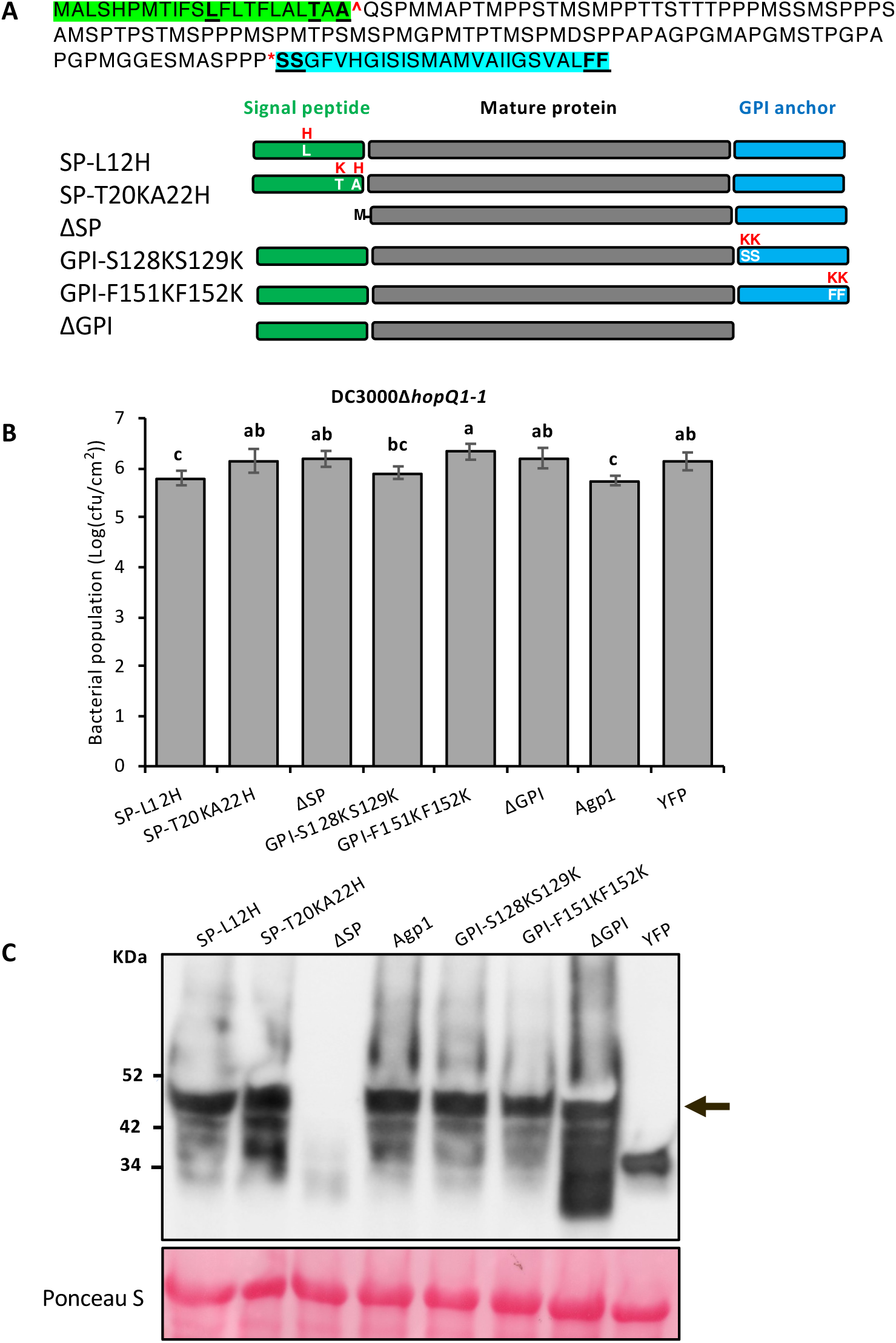
The signal peptide (SP) and glycosylphosphatidylinositol (GPI) anchor play a role in Agp1-mediated immunity to *Pst*. **A**, Top: amino acid sequence of the Agp1 protein. Signal peptide sequence is highlighted in green and the GPI-anchored sequence is highlighted in blue. Schematics show the substituted amino acids or deletions of the Agp1 protein, each fused to an HA epitope tag. **B**, Leaves of five-week-old *N. benthamiana* plants were syringe-infiltrated with *Agrobacterium* (1D1249) strains (OD = 0.5) carrying a binary expression vector expressing each gene. Two days later, the same agroinfiltrated spots were syringe-infiltrated with 5 × 10^4^ cfu/mL DC3000*ΔhopQ1-1. Pst* DC3000 populations were measured two days after the second infiltration. Bars show means ± SD. Different letters indicate significant differences based on a one-way ANOVA followed by Student’s *t* test (p < 0.05). Three or four plants were tested with each gene in each experiment. The experiments were performed twice with similar results. **C**, Proteins were extracted from *N. benthamiana* leaves expressing each Agp1:HA variant two days after agroinfiltration. Proteins were detected by immunoblotting with an α-HA antibody.

### Loss of Nrd1 function has no effect on the transcript abundance of multiple PTI-associated genes

Multiple tomato immunity-associated genes including *Bti9, Core, Fls2, Fls3* and *Wak1* play important roles in PTI responses (Hind et al., 2016; Roberts et al., 2020; Rosli et al., 2013; Wang et al., 2016; Zeng et al., 2012; Zheng et al., 2019). We analyzed our RNA-Seq data to determine whether loss of Nrd1 function affects transcript abundance of these immunity-associated genes upon inoculation with the PTI-inducing strain DC3000Δ*avrPto*Δ*avrPtoB* (**Supplemental Table 3**). Interestingly, no significant difference of the transcript abundance of these genes was observed between the two Δnrd1 mutant lines and the wild-type plants. It has been reported that *Bti9, Core, Fls2, Fls3* and *Wak1* are greatly upregulated in wild-type RG-PtoR plants upon inoculation with DC3000Δ*avrPto*Δ*avrPtoB* (Rosli et al., 2013), indicating that these genes were also induced in the Δnrd1 mutants by this *Pst* strain regardless of the loss of Nrd1function. In all, the enhanced immunity observed in the Δnrd1 mutants is likely due to the activation of key components of PRR signaling (*Fls2*/*Fls3*/*Wak1*, etc.) as well as loss of Nrd1-regulated suppression of the defense gene *Agp1*.

## Discussion

The *Nrd1* gene was originally identified from a small subset of 44 genes whose transcript abundance in tomato leaves increased in response to flgII-28 but not in response to flg22 or csp22 (Rosli et al., 2013). This specificity was subsequently confirmed by RT-qPCR and *Nrd1* is therefore useful as a reporter gene for the Fls3 pathway (Roberts et al., 2020). Because the gene is induced by flgII-28 we anticipated that a loss-of-function mutation in *Nrd1* might lead to loss of certain aspects of pattern-triggered immunity. However, unexpectedly, two independent Δnrd1 mutants showed enhanced resistance specifically to *Pst* DC3000, indicating the Nrd1 protein acts as a negative regulator of resistance to this *Pst* strain. An RNA-Seq analysis of the Δnrd1 mutants identified a small number of genes whose transcript abundance is either increased or decreased in an Nrd1-dependent manner and we hypothesized these genes might play a role in defense or susceptibility, respectively. Overexpression of one of the putative defense genes, *Agp1*, encoding an arabinogalactan protein, did in fact enhance resistance to DC3000, suggesting that it plays a role in the enhanced resistance of the Δnrd1 mutants. Here we place Nrd1 in the context of previous reports of negative regulators of immunity, and we discuss the possible role of Agp1 in defense, propose a model for Fls3-specific transcriptional reprogramming, and consider the prospect that Nrd1/Agp1 might be used to identify a unique component of *Pst* DC3000 that is involved in the enhanced resistance observed in the Δnrd1 mutants.

Negative regulators of plant immunity can be viewed as susceptibility (S) genes since their expression allows enhanced growth of the pathogen and accordingly enhanced disease (van Schie and Takken, 2014). S genes have been classified into those that play a role in host recognition, suppression of host defenses, or in pathogen sustenance and they encode diverse proteins including transporters, protein kinases, membrane-associated proteins (e.g., Mlo), and enzymes (e.g., Dmr6) (Santillan Martinez et al., 2020; Thomazella et al., 2021; van Schie and Takken, 2014; Zheng et al., 2013). Of particular relevance here, several S genes encode transcription factors in the bHLH, bZIP, ERF, and WRKY families (Fan et al., 2014; Fang et al., 2021; Jin et al., 2011; Lu et al., 2020; Prior et al., 2021; Schwartz et al., 2017; Wang et al., 2015a). Similar to Nrd1, a few bHLH transcription factors have been found previously to act as negative regulators of disease resistance in plants. For instance, two tomato bHLH genes, *SlbHLH3* and *SlbHLH6*, are up-regulated by the transcription activator-like effector (TALE) AvrHah1 in *Xanthomonas gardneri* and promote susceptibility of tomato to bacterial spot disease (Schwartz et al., 2017). bHLH TFs in other plant species, including the well-characterized HBI1 in *Arabidopsis thaliana*, negatively regulates a subset of genes involved in plant immunity and mediates a trade-off between growth and immunity in plants (Fan et al., 2014). In contrast to these bHLH negative regulators which are either induced by bacterial effectors (Schwartz et al., 2017) or suppressed by MAMPs or other bacterial components (Fan et al., 2014), the *Nrd1* gene is induced specifically by a flagellin-derived MAMP flgII-28 but acts in a way that promotes bacterial pathogenesis.

The tomato receptor Fls3 binds flgII-28 and works in concert with the co-receptor BAK1 (in tomato, Serk3A and/or Serk3B) to activate intracellular signaling (Hind et al., 2016). Our present and previous RNA-Seq analysis and the phenotype of the Δnrd1 mutants together are consistent with a model in which Fls3 activates both resistance-enhancing and susceptibility-enhancing responses (**Fig. 6**). To resist *Pst* infection, Fls3 and other PRRs activate PTI responses leading to the rapid generation of ROS, activation of MAPKs and extensive changes in transcriptional programming that inhibit *Pst* growth. Fls3 also induces *Nrd1* gene expression and likely increases Nrd1 protein abundance which, we propose, suppresses a subset of defense genes and induces a subset of susceptibility genes promoting tomato susceptibility to *Pst* infection. In a loss-of-function mutation in *Nrd1* the subset of defense genes, including *Agp1*, are no longer suppressed (or are induced) and S genes are not expressed, leading to enhanced *Pst* resistance. Additionally, in the Δnrd1 mutants, multiple well-characterized defense genes including *Bti9, Core, Fls2, Fls3* and *Wak1* are still induced upon *Pst* inoculation (**Supplemental Table 3**), and ROS production and MAPK activation are not compromised (**Fig. 3**), suggesting that the observed increased resistance in the Δnrd1 mutants is due to the activation of key PRR signaling components as well as loss of Nrd1-regulated suppression of some defense genes such as *Agp1* and/or loss of Nrd1-regulated induction of certain S genes.

**Figure 6.**
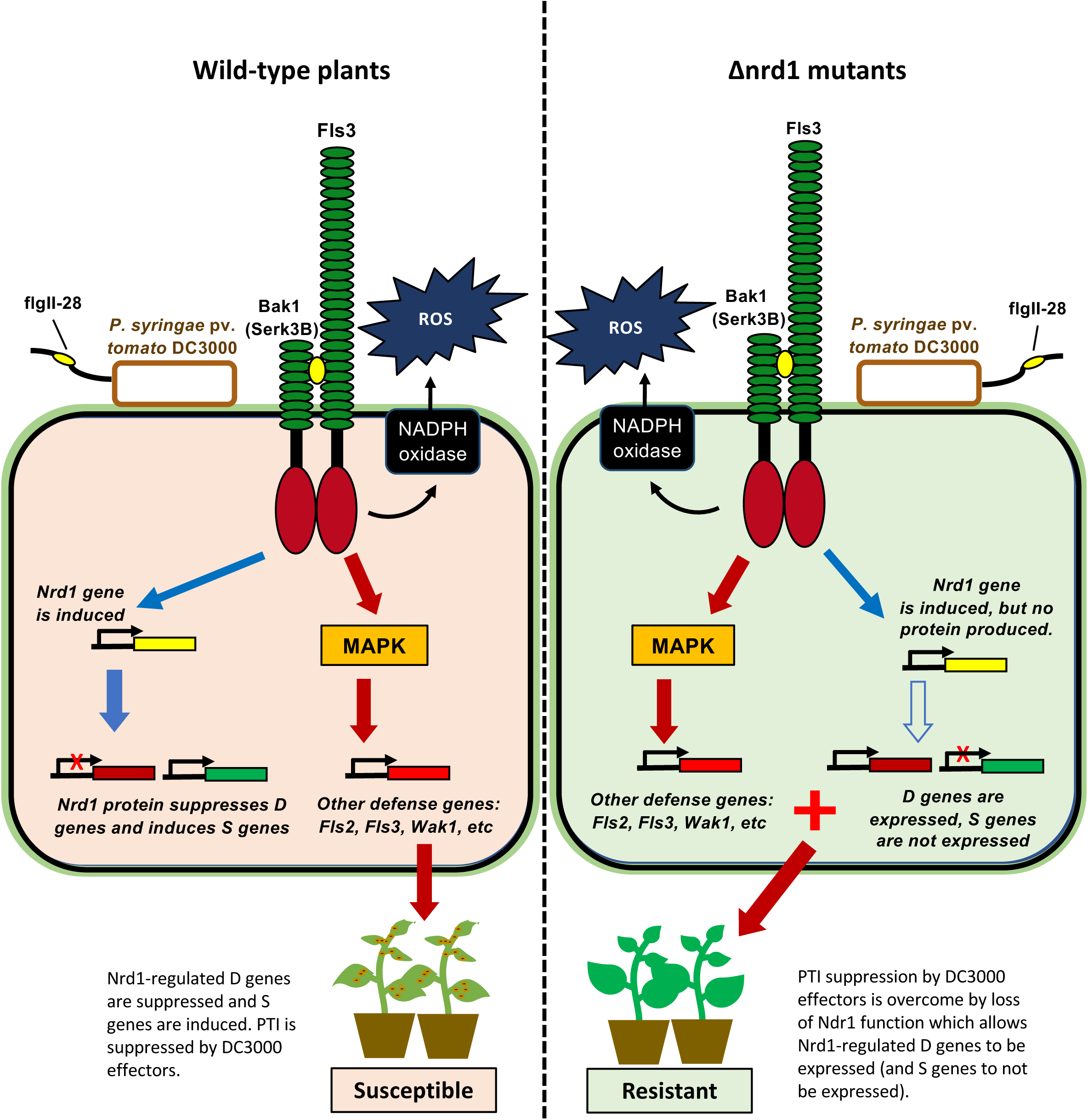
Proposed model for the enhanced resistance seen in Δnrd1 mutants. Fls3 appears to regulate two opposing host responses: 1) To resist *Pst* infection, Fls3 and other PRRs induce ROS, MAPK and other defense responses which inhibit *Pst* growth. 2) Fls3 also induces Nrd1 gene expression, and increases Nrd1 protein abundance, which suppresses a subset of defense genes and also induces a subset of susceptibility genes further promoting susceptibility to *Pst* infection. When Nrd1 is mutated, the subset of defense genes, including Agp1, are no longer suppressed (or are induced) and S genes are not expressed leading to enhanced resistance to *Pst*.

The discovery that overexpression of the tomato *Agp1* gene significantly reduced DC3000 populations in leaves further reinforces the importance of the plant cell wall as the location for key immunity-associated activities (Bacete et al., 2018; Molina et al., 2021). Arabinogalactan-proteins (AGPs) belong to a large family of cell wall hydroxyproline-rich glycoproteins that are involved in diverse biological processes including plant growth and development and plant-microbe interactions (Gaspar et al., 2004; Seifert and Roberts, 2007). Classical AGPs contain an N-terminal hydrophobic secretion signal, a central domain rich in PAST (Pro, Ala, Ser, and Thr) residues, and a hydrophobic C-terminal sequence that directs the attachment of GPI anchor (Silva et al., 2020), whose presence or absence has been demonstrated to play a major impact on the host immune response to pathogen infection (Butikofer et al., 2001). GPI modification also allows the defense-associated protein NDR1 to attach on the outer surface of the plasma membrane, thus positively regulating disease resistance to multiple bacterial and fungal pathogens (Century et al., 1997; Century et al., 1995; Coppinger et al., 2004). In yeast, lesions in GPI-anchor production prevent certain proteins reaching the cell surface leading to cell wall defects and even death (Kinoshita et al., 1997). Consistent with this, we found removal of the GPI anchor from Agp1 caused a loss of *N. benthamiana* resistance to *Pst* DC3000, indicating the essential role of GPI anchor on Agp1 function in the tomato immune response, likely by disrupting the association of the Agp1 protein with the extracellular face of the plasma membrane. Additionally, Agp1 appeared to be heavily glycosylated, a common post-translational modification in AGPs that might regulate protein conformation, activity and stability in host-pathogen interactions (Lin et al., 2020).

The molecular mechanisms of AGPs in plant-microbe interactions remain largely unknown. It has been proposed that GPI-anchored proteins can be involved in signaling via phospholipase cleavage of the protein from the lipid anchor or via interactions with other plasma membrane or cell wall-associated proteins that are able to activate signaling pathways (Schultz et al., 1998; Schultz and Harrison, 2008; Yeats et al., 2018; Zhou, 2019). It is intriguing to speculate that GPI-anchored Agp1 might act in a complex with PRRs and modulate ligand recognition specificity (Yeats et al., 2018; Zhou, 2019) or that Agp1 interacts with the cell-wall associated kinase *SlWak1* (Zhang et al., 2020) after release of Agp1 from the plasma membrane by cleavage of the GPI anchor; AGP epitopes have been reported to co-localize with Waks in tobacco protoplasts (Gens et al., 2000). Degradation products of AGPs might also function as damage-associated molecular patterns (DAMPs) eliciting a defense response (Villa-Rivera et al., 2021). In this regard, Arabidopsis WAK1 has been demonstrated to be a receptor of oligogalacturonides (OGs), an important component of some DAMPs (Brutus et al., 2010). The observation that AGPs localize in lipid rafts where many receptor proteins are clustered further supports the hypothesis that Agp1 might associate with certain defense-associated receptors (Ellis et al., 2010). Although these studies suggest possible molecular mechanisms of AGPs in plant-microbe interaction, more experiments are needed to understand how Agp1 enhances plant defense.

Loss-of-function mutations in S genes might offer a promising approach to enhancing broad-spectrum disease resistance, as long as the mutation does not have pleiotropic detrimental effects. There are several examples of this strategy in the literature, although none yet involve a bHLH transcription factor (Hanika et al., 2021; Santillan Martinez et al., 2020; Seifert and Roberts, 2007; Sun et al., 2016; Thomazella et al., 2021; van Schie and Takken, 2014; Zheng et al., 2013). In contrast to such broad-spectrum activity, the enhanced resistance in the Δnrd1 mutants appears specific to *Pst* DC3000 as the Δnrd1 mutants were susceptible to four other strains of *Pst* and to the bacterial pathogen *Xanthomonas* (**Supplemental Table 1**). In light of this, although we saw no detrimental morphological or growth defects in the Δnrd1 mutants they will likely not be generally useful for controlling bacterial speck disease. However, our results do raise the possibility that DC3000 expresses a unique component, lacking in other *Pst* strains, that is recognized by the Δnrd1 mutants. The future identification of such a *Pst* component might lead to the discovery of a novel host recognition mechanism.

## Materials and Methods

### Generation of *Nrd1* tomato mutants using CRISPR/Cas9

To generate the Δnrd1 mutants in the tomato cultivar Rio Grande (RG)-PtoR, which has the *Pto* and *Prf* genes, we designed a guide RNA (5’-GTAGTCCAGAAAAGCTAGAC-3’) that targets the first exon of *Nrd1* using the software Geneious R11 (Kearse et al., 2012). The gRNA cassette was cloned into the p201N:Cas9 binary vector as described previously (Jacobs et al., 2017). Tomato transformation was performed at the Biotechnology Center at the Boyce Thompson Institute as described previously (Zhang et al., 2020). Mutations were confirmed by Sanger sequencing at the Biotechnology Resource Center (BRC) at Cornell University.

### Phylogenetic analyses

The Nrd1 protein sequence was used as a query sequence to search for related sequences in tomato, *Arabidopsis*, and rice using the NCBI BLAST (https://blast.ncbi.nlm.nih.gov/Blast.cgi). Amino acid alignments were performed by ClustalW (https://www.genome.jp/tools-bin/clustalw). Phylogenetic trees were constructed with MEGA-X (Kumar et al., 2018) using the maximum likelihood method and JTT matrix-based model (Jones et al., 1992). Bootstrap analysis with 1000 replicates was performed. Positions containing gaps and missing data were eliminated.

### Bacterial inoculation

Four-week-old Δnrd1 and wild-type plants were vacuum-infiltrated with various *Pst* DC3000 strains at different titers, including DC3000Δ*avrPto*Δ*avrPtoB* (DC3000ΔΔ) or DC3000Δ*avrPto*Δ*avrPtoB*Δ*fliC* (DC3000ΔΔΔ) at 5 × 10^4^ cfu/mL or DC3000 at 1 × 10^6^ cfu/mL. Bacterial populations were measured at 3 h (Day 0) and two days after inoculation (Day 2). Photographs of disease symptoms were taken five or six days after bacterial inoculation.

### ROS assay

ROS production was measured as described previously (Clarke et al., 2013). In brief, leaf discs were collected and floated in water overnight. Water was then removed and replaced with a solution containing flg22 (QRLSTGSRINSAKDDAAGLQIA) or flgII-28 (ESTNILQRMRELAVQSRNDSNSSTDRDA) at the indicated concentrations, in combination with 34 µg/mL luminol (Sigma-Aldrich) and 20 µg/mL horseradish peroxidase. ROS production was measured using a Synergy 2 microplate reader (BioTek).

### MAPK phosphorylation assay

MAPK phosphorylation assay was performed as described previously (Zhang et al., 2020). Six leaf discs of Δnrd1 mutant and wild-type plants were floated in water overnight. The leaf discs were then incubated with flg22 or flgII-28 at desired concentrations, or water only for 10 min, and immediately frozen in liquid nitrogen. Protein was extracted using buffer containing 50 mM Tris-HCl (pH 7.5), 10% glycerol, 2 mM EDTA, 1% Triton X-100, 5 mM DTT, 1% protease inhibitor cocktail (Sigma-Aldrich), 0.5% Phosphatase inhibitor cocktail 2 (Sigma-Aldrich). MAPK phosphorylation was determined using an anti-phospho-p44/42 MAPK(Erk1/2) antibody (anti-pMAPK; Cell Signaling).

### Construct generation

The coding region of each putative defense or susceptibility gene was amplified from tomato cDNA using Phusion Hot Start II DNA polymerase (ThermoFisher Scientific) and gene-specific primers (**Supplemental Table 4**), then cloned into pJLSmart (Mathieu et al., 2014) by Gibson assembly. The gene expression cassette in pJLSmart was then cloned into the destiny vector pGWB414 via recombination reactions using LR Clonase II (ThermoFisher Scientific). Vectors were confirmed by Sanger sequencing and then transformed into *Agrobacterium* strain 1D1249 for transient expression and agromonas assay in *N. benthamiana*.

Amino acid substitutions in the signal peptide and GPI-anchor sequences of the Agp1 protein were determined using SignalP-5.0 (Almagro Armenteros et al., 2019) and NetGPI-1.1(Gíslason et al., 2021). Amino acid substitutions were generated with the Q5 site-directed mutagenesis kit (NEB) with specific primers (**Supplemental Table 4**). The signal peptide sequence (retaining ATG) and the GPI sequence were deleted by PCR with specific primers using Phusion Hot Start II DNA polymerase (**Supplemental Table 4**). All mutated fragments were first cloned into pJLSmart by Gibson assembly and then pGWB414 by LR reaction.

### Agromonas assay

The agromonas assays were performed as described (Buscaill et al., 2021). Briefly, *Agrobacterium* strains 1D1249 carrying a binary vector (pGWB414) expressing the gene of interest was syringe-infiltrated into leaves of four-week-old *N. benthamiana* plants. Two days later, the same agroinfiltrated spots were syringe-infiltrated with either DC3000Δ*hopQ1-1* or DC3000Δ*hopQ1*-*1*Δ*avrPto*Δ*avrPtoB* at 5 × 10^4^ cfu/mL. Bacterial populations were measured by serial dilutions on LB medium supplemented with 10 µg/ml cetrimide, 10 µg/ml fucidin and 50 µg/ml cephaloridine (CFC; Oxoid™ C-F-C Supplement) two days after *Pst* inoculation.

### Immunoblotting

Total protein was extracted from *N. benthamiana* leaves using 250 µl extraction buffer consisting of 62.5 mM Tris-HCl (pH 6.8), 2% SDS (v/v), 10% glycerol and 5% β-mercaptoethanol. A 12 µL soluble protein solution mixed with 4X Laemmli sample buffer were boiled at 95°C for 5 min before loaded for gel electrophoresis. Protein was loaded on 4% - 20% precast SDS-PAGE gel (Bio-Rad), blotted on PVDF membrane (Merck Millipore), inoculated with α-HA primary antibody (1:7000; v/v) and α-rat-HRP secondary antibody (1:10000; v/v), and developed with Piece ECL plus substrate (Thermo Scientific) for 5 min.

### RNA-Seq

Five-week-old wild-type RG-PtoR and the two lines of Δnrd1 mutants were vacuum infiltrated with a suspension of DC3000Δ*avrPto*Δ*avrPtoB* at 5 × 10^6^ cfu/mL. Four biological replicates were performed for each treatment. Tissue samples were collected at 6 h after infiltration. Total RNA was isolated with the RNeasy Plant Mini Kit (Qiagen) according to the manufacturer’s instructions. RNA was treated with DNase by column-based purification (RNase-Free DNase Kit, Qiagen). RNA libraries were prepared and sequenced on an Illumina HiSeq 4000 system. Raw RNA-Seq reads were processed to remove adaptors and low-quality sequences using Trimmomatic (version 0.36) with default parameters (Bolger et al., 2014). The remaining cleaned reads were aligned to the ribosomal RNA database (Quast et al., 2013) using bowtie (version 1.1.2; (Langmead, 2010)) allowing up to three mismatches, and those aligned were discarded. The remaining cleaned reads were mapped to the tomato reference genome (SL4.0 and ITAG4.1) using HISAT2 (version 2.1.0; (Kim et al., 2019)) with default parameters. Based on the alignments, raw read counts for each gene were calculated using HTSeq-count (Anders et al., 2015) and normalized to fragments per kilobase of transcript per million mapped fragments (FPKM). Raw read counts were then fed to DESeq2 (Love et al., 2014) to identify differentially expressed genes (DEGs), with a cutoff of adjusted *P* value < 0.05 and fold change > 2.

## Acknowledgments

We thank Liam Cleary, Brian Bell, Jay Miller, and Joe Ettenberger for plant care, and Joyce Van Eck for tomato transformation. Funding was provided by National Science Foundation grant IOS-1546625 (GBM and ZF).

## Author contributions

GBM and NZ conceived and designed the experiments. NZ designed gRNAs, constructed vectors, performed genotyping and phenotyping experiments, and analyzed the data. CH performed ROS assays. ZF and XS analyzed RNA-Seq data. NZ and GBM interpreted the data and wrote the manuscript. All the authors read and approved the manuscript.

## Supplemental Information

**Supplemental Figure 1.**
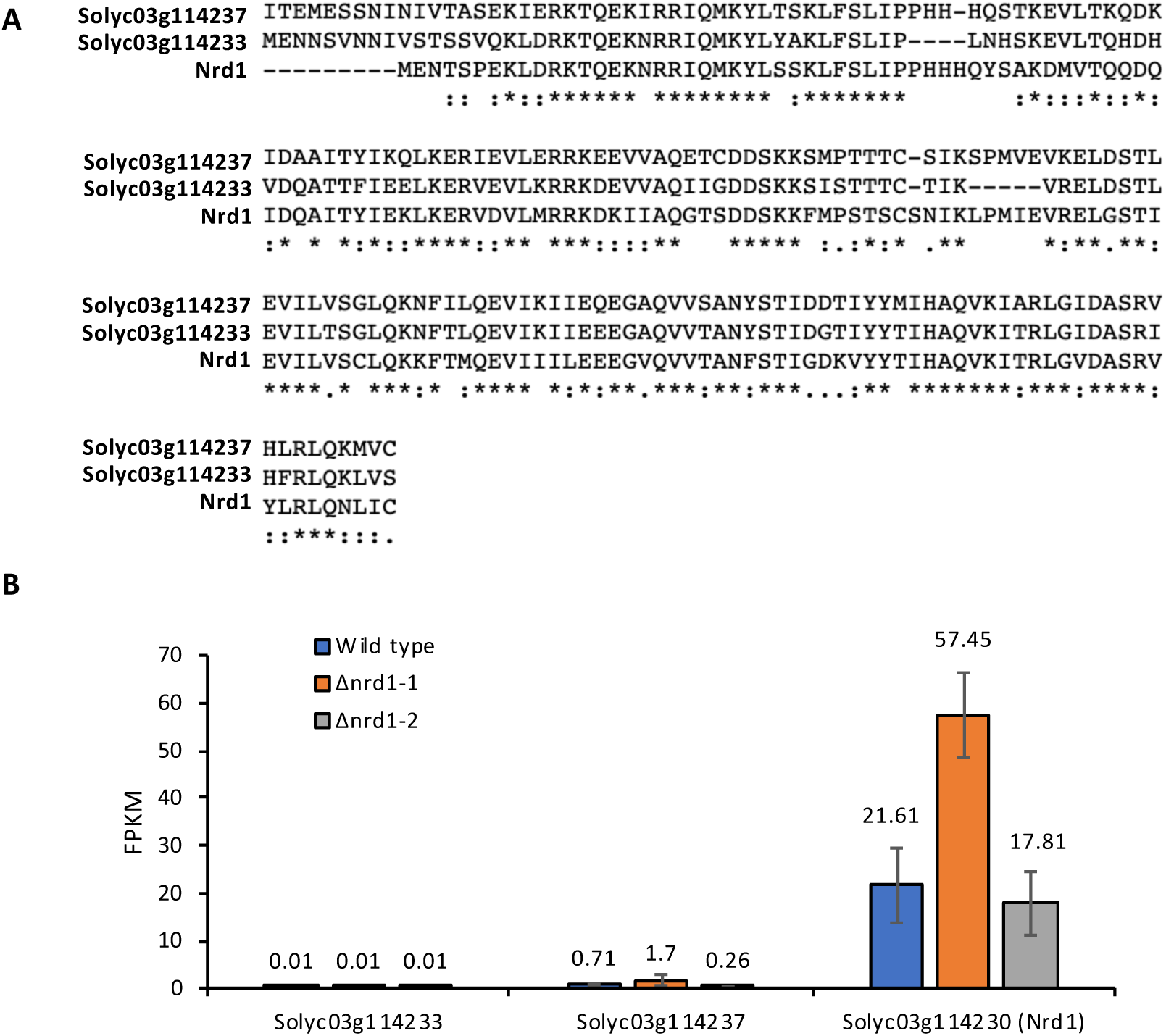
The two closest Nrd1 homologs in tomato. **A**, Amino acid sequence alignment of the two closest tomato Nrd1 homologs. **B**, Transcript abundance of the two closest tomato Nrd1 homologs in wildtype and Δnrd1 mutants, 6 h after treatment with 5 × 10^6^ cfu/mL DC3000Δ*avrPto*Δ*avrPtoB* (DC3000ΔΔ).

**Supplemental Figure 2.**
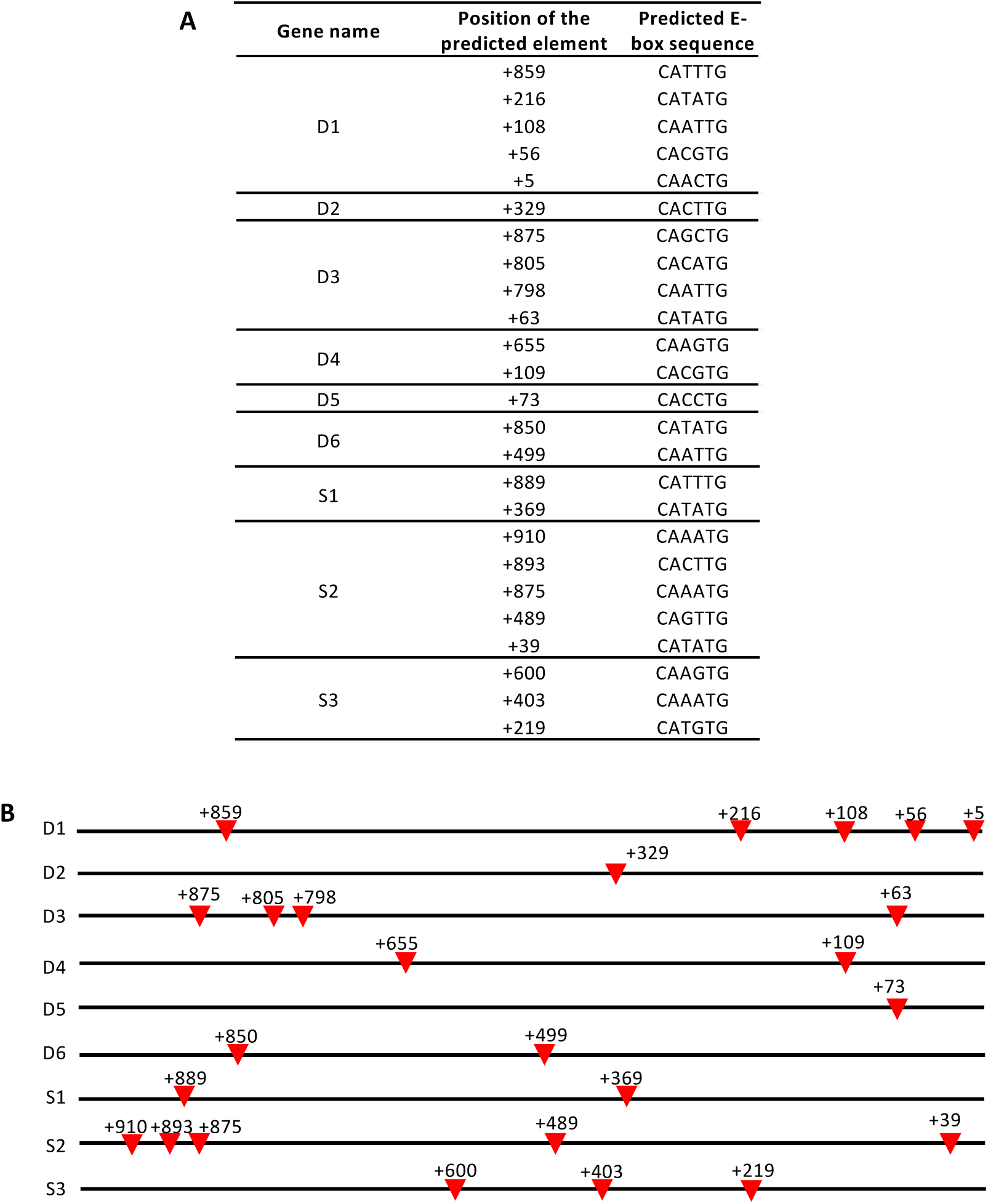
Predicted E-box elements (CANNTG) in Nrd1-regulated putative defense and susceptibility genes. **A**-**B**, 1 kb DNA sequence upstream of the 5’ untranslated region (5’UTR) or coding region (CDS) of each gene was analyzed by PlantPan2.0 (Chow et al., 2016) to identify potential bHLH-binding elements, denoted with inverted red triangles. The first nucleotide of the promoter sequence upstream of 5’UTR or CDS is designated as “+1”.

**Supplemental Figure 3.**
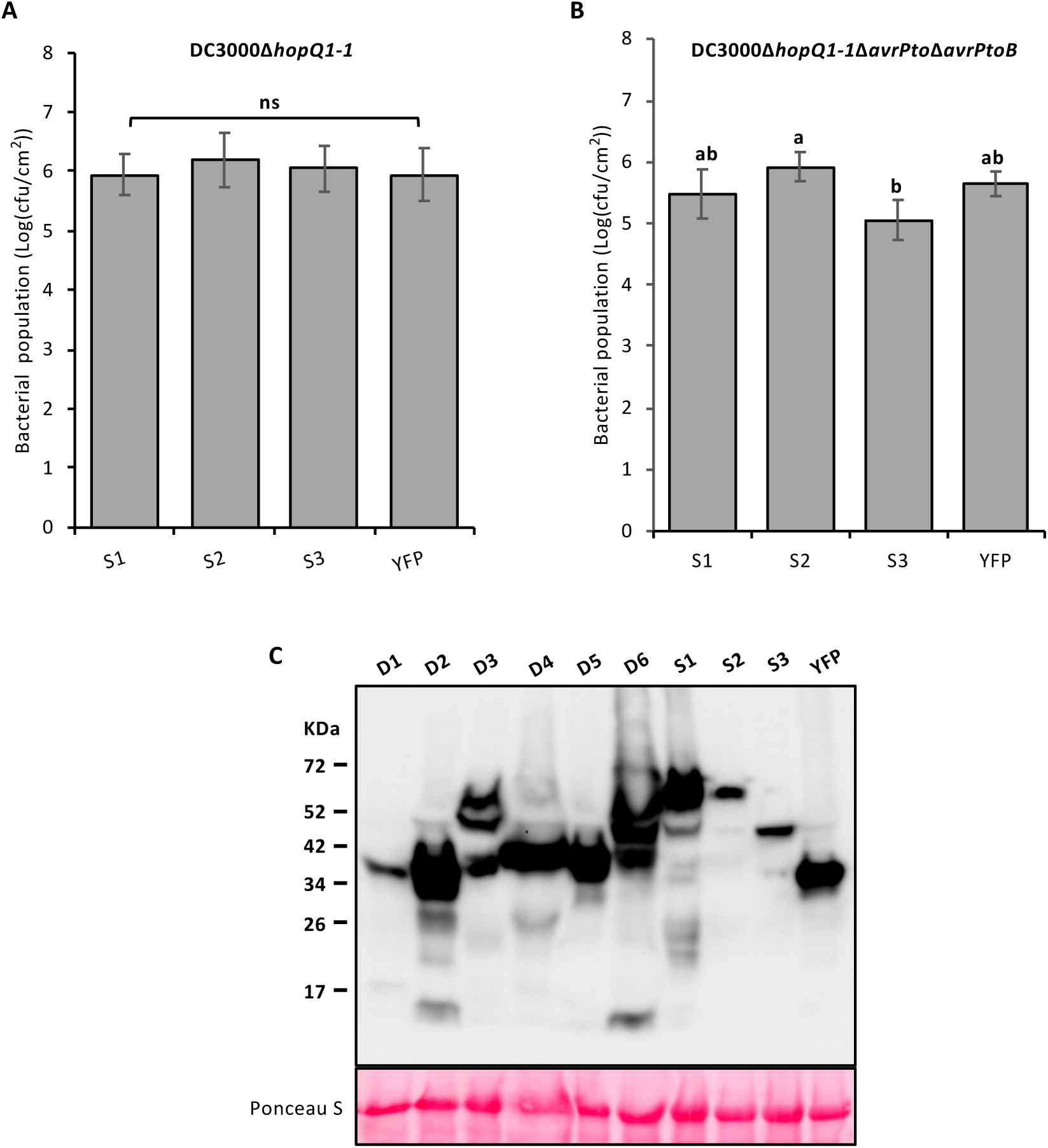
Testing putative susceptibility genes in an agromonas assay and confirming expression of the D and S proteins. **A**-**B**, Five-week-old *Nicotiana benthamiana* plants were syringe-infiltrated with *Agrobacterium* strains (1D1249) containing a binary expression vector expressing each gene (OD = 0.5). Two days later, the same agro-infiltrated spots were syringe-infiltrated with 5 × 10^4^ cfu/mL DC3000*ΔhopQ1-1* (**A**) or 5 × 10^4^ cfu/mL DC3000*ΔhopQ1-1*Δ*avrPto*Δ*avrPtoB* (**B**). Bacterial populations were measured two days after infiltration. **A**-**B**, Bars represent means ± SD. Different letters indicate significant differences based on a one-way ANOVA followed by Student’s *t* test (p < 0.05). ns, no significant difference. **C**, Protein expression by western blotting. Proteins were extracted from *N. benthamiana* leaves two days after agroinfiltration. S proteins with an HA epitope tag were detected by immunoblotting with α-HA antibody.

**Supplemental Figure 4.**
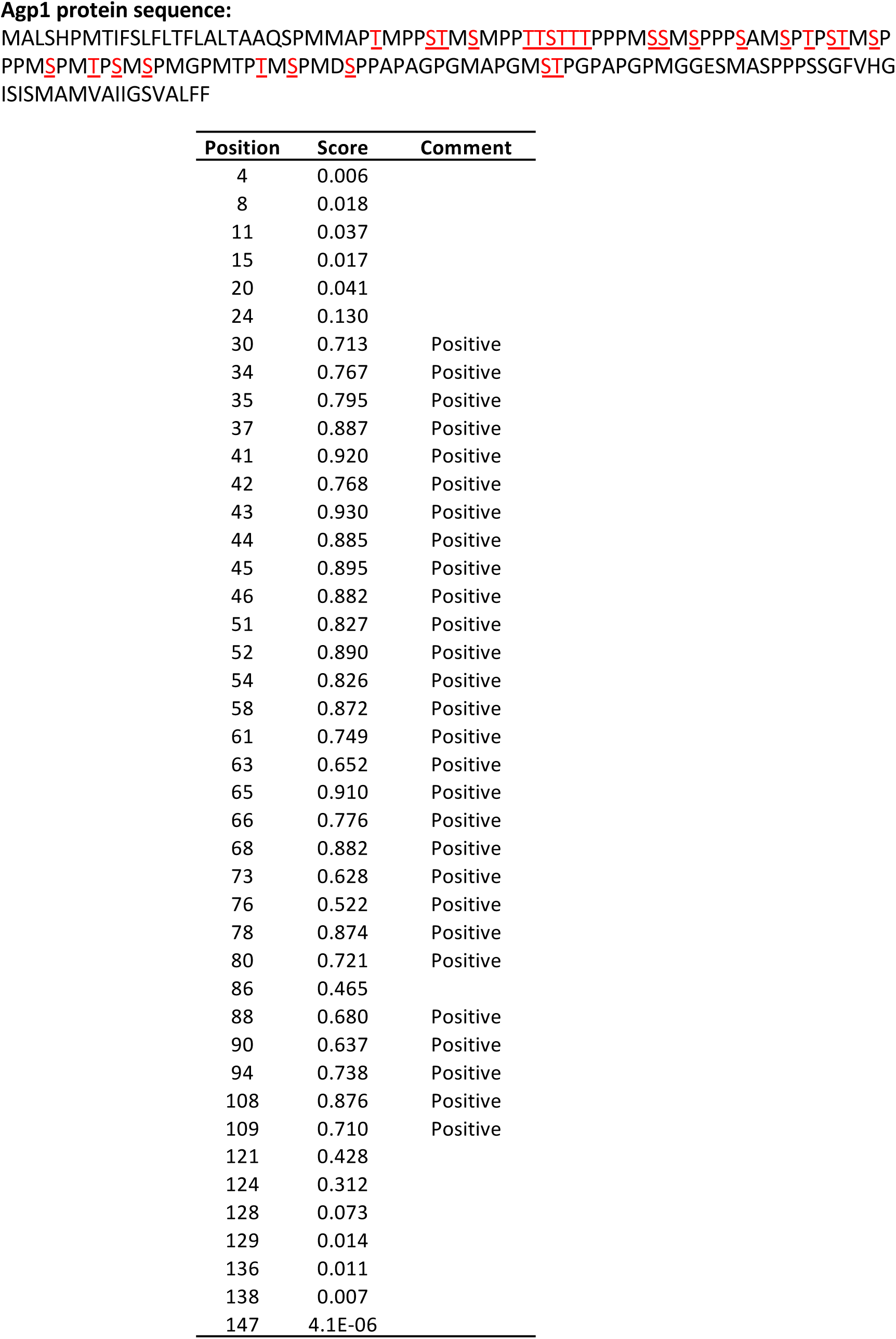
Prediction of glycosylation sites in the Agp1 protein. Glycosylation sites were predicted using NetOGlyc-4.0 (Steentoft et al., 2013) with a cutoff score higher than 0.5 (marked as “Positive”). The first amino acid of Agp1 protein is designated as position “1”. Positive sites of glycosylated amino acids are highlighted in red and underlined.

**Supplemental Table 1.**
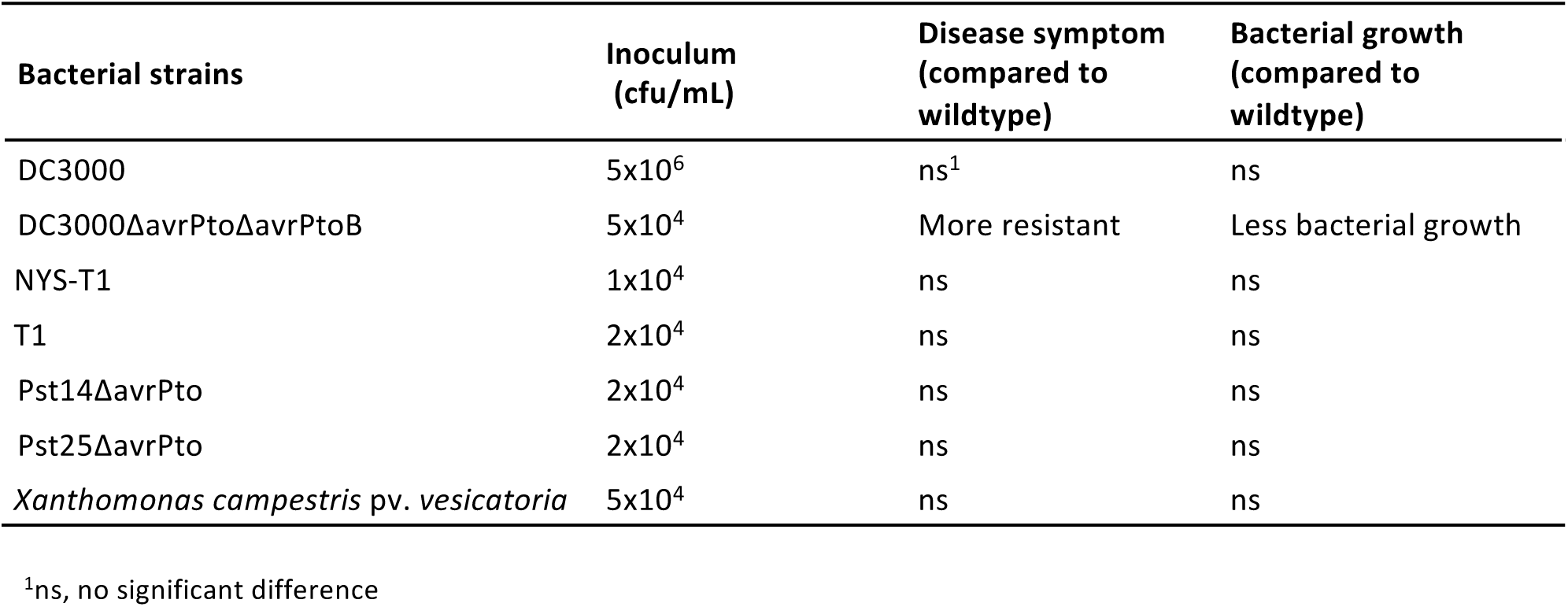
Summary of disease assays with the Δnrd1 mutant plants.

**Supplemental Table 2.**
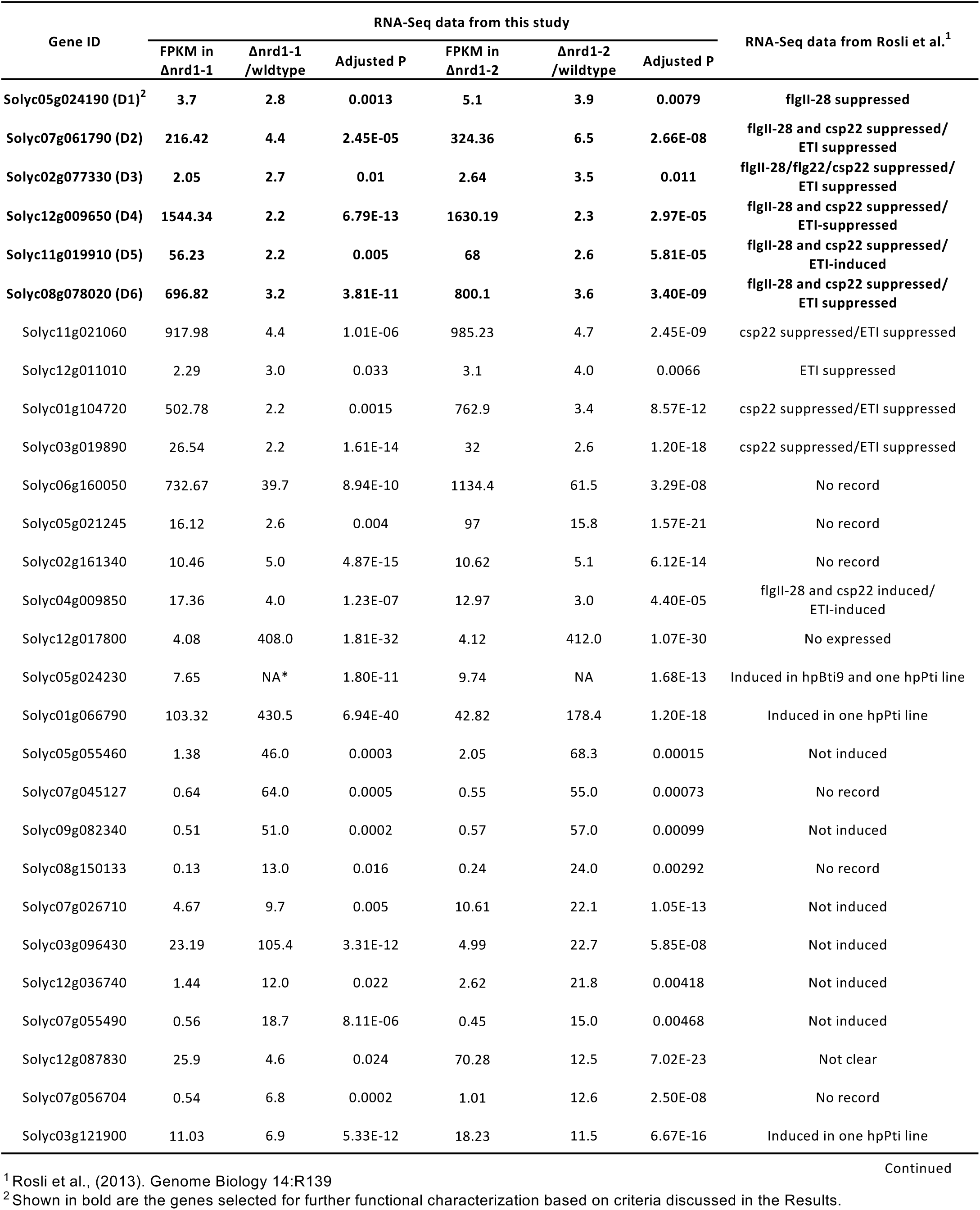
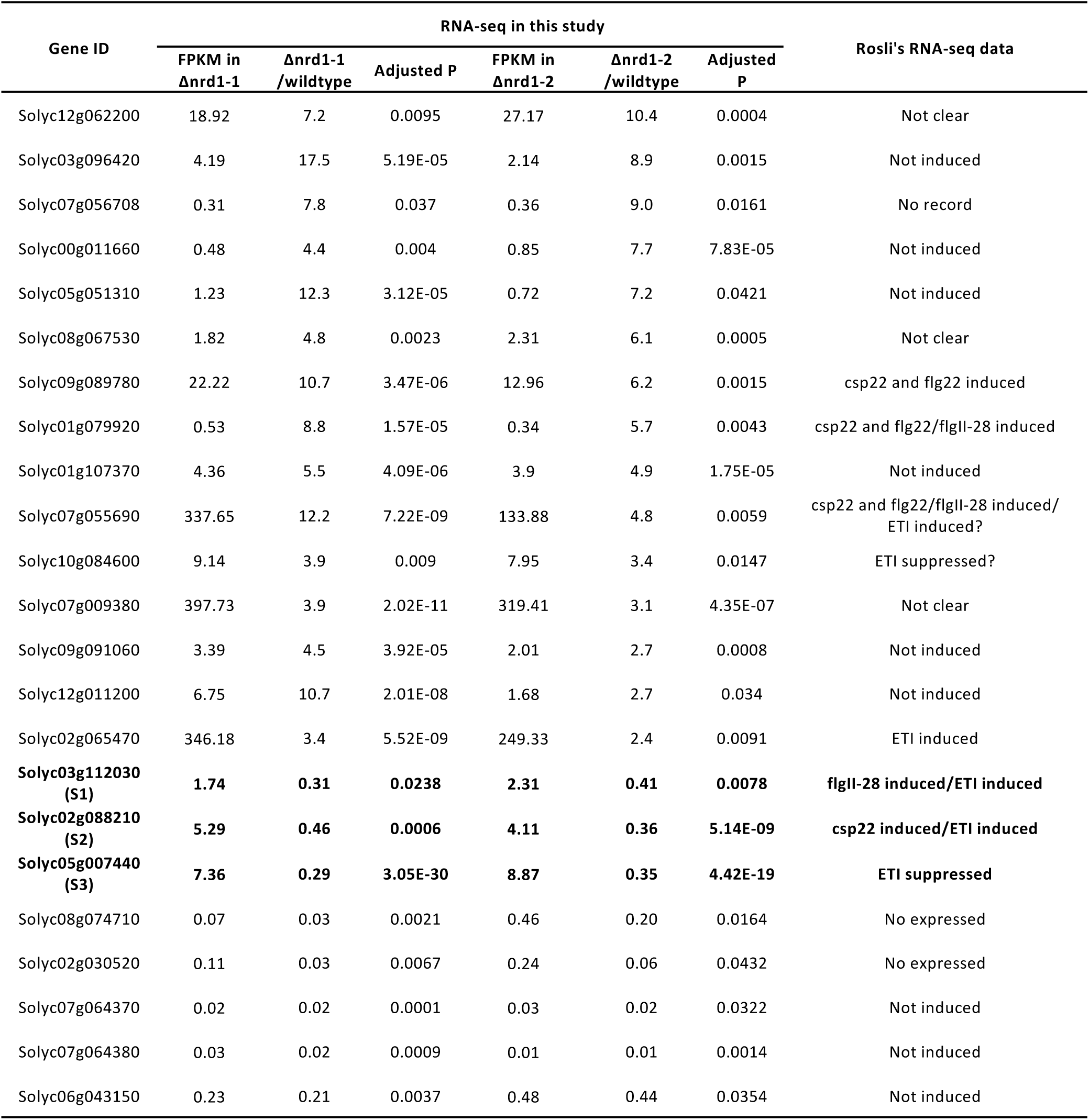
The 51 *Nrd1*-regulated putative defense and susceptibility genes identified by RNA-Seq.

**Supplemental Table 3.** RNA-Seq data showing the transcript abundance of selected immunity-associated genes in Δnrd1 mutants when inoculated with DC3000Δ*avrPto*Δ*avrPtoB*.

| Gene name | Gene ID | FPKM | | $\Delta nrd1-1$ /wildtype | Adjusted P | FPKM | | $\Delta nrd1-2$ /wildtype | Adjusted P |
| --- | --- | --- | --- | --- | --- | --- | --- | --- | --- |
| | | $\Delta nrd1-1$ | Wildtype | | | $\Delta nrd1-2$ | Wildtype | | |
| Wak1 | Solyc09g014720 | 15.32 | 10.61 | 1.44 | 0.45 | 8.1 | 10.61 | 0.76 | 0.75 |
| Fls2.1 | Solyc02g070890 | 4.63 | 3.03 | 1.53 | 0.03 | 3.37 | 3.03 | 1.11 | 0.74 |
| Fls3 | Solyc04g009640 | 16.93 | 10.98 | 1.54 | 0.37 | 7.66 | 10.98 | 0.70 | 0.46 |
| Bti9 | Solyc07g049180 | 31.78 | 33.9 | 0.94 | 0.99 | 26.18 | 33.9 | 0.77 | 0.01 |
| Core | Solyc03g096190 | 2.05 | 1.3 | 1.58 | 0.52 | 0.81 | 1.3 | 0.62 | 0.27 |

**Supplemental Table 4.**
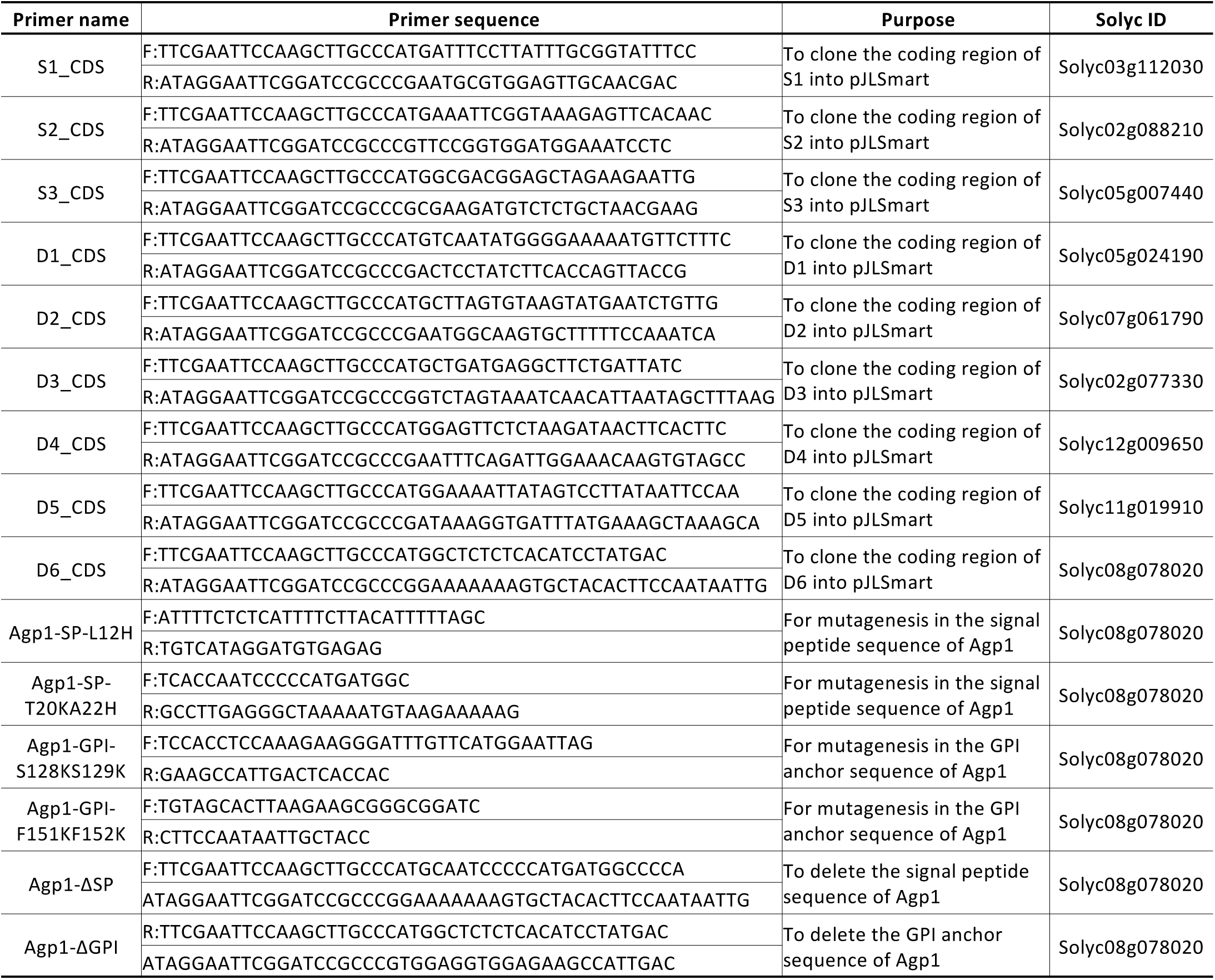
Primers used in this study.

